# *Brucella abortus* induces a Warburg shift in host metabolism that is linked to enhanced intracellular survival of the pathogen

**DOI:** 10.1101/120527

**Authors:** Daniel M. Czyż, Jonathan Willett, Sean Crosson

## Abstract

Intracellular bacterial pathogens exploit host cell resources to replicate and survive inside the host. Targeting these host systems is one promising approach to developing novel antimicrobials to treat intracellular infections. We show that human macrophage-like cells infected with *Brucella abortus* undergo a metabolic shift characterized by attenuated tricarboxylic acid cycle metabolism, reduced amino acid consumption, altered mitochondrial localization, and increased lactate production. This shift to an aerobic glycolytic state resembles the Warburg effect, a change in energy production that is well-described in cancer cells, and also occurs in activated inflammatory cells. *B. abortus* efficiently uses lactic acid as its sole carbon and energy source and requires the ability to metabolize lactate for normal survival in human macrophage-like cells. We demonstrate that chemical inhibitors of host glycolysis and lactate production do not affect *in vitro* growth of *B. abortus* in axenic culture, but decrease its survival in the intracellular niche. Our data support a model in which infection shifts host metabolism to a Warburg-like state, and *B. abortus* uses this change in metabolism to promote intracellular survival. Pharmacological perturbation of these features of host cell metabolism may be a useful strategy to inhibit infection by intracellular pathogens.

**IMPORTANCE:** *Brucella* spp. are intracellular bacterial pathogens that cause disease in a range of mammals, including livestock. Transmission from livestock to humans is common and can lead to chronic human disease. Human macrophage-like cells infected with *Brucella abortus* undergo a Warburg-like metabolic shift to an aerobic glycolytic state where the host cells produce lactic acid and have reduced amino acid catabolism. We provide evidence that the pathogen can exploit this change in host metabolism to support growth and survival in the intracellular niche. Drugs that inhibit this shift in host cell metabolism inhibit intracellular replication and decrease the survival of *B. abortus* in an *in vitro* infection model; these drugs may be broadly useful therapeutics for intracellular infections.

## INTRODUCTION

Macrophages can be activated by pathogen-associated pro-inflammatory molecules such as lipopolysaccharide (LPS). Classically activated (i.e. M1) macrophages undergo a major metabolic shift in which the tricarboxylic acid (TCA) cycle is downregulated, and energy production transitions from oxidative phosphorylation to the less efficient process of aerobic glycolysis (1, 2). This change in metabolism, known as the Warburg effect (3), is a well-described metabolic feature of cancer cells (4). Increased glucose consumption through glycolysis is thought to increase adenosine triphosphate (ATP) levels; a corresponding increase in pentose phosphate pathway activity generates nicotinamide adenine dinucleotide phosphate (NADPH), which is subsequently used by NADPH oxidases to generate reactive oxygen species (ROS) (5). This metabolic shift enables cells to launch general antimicrobial defenses by increasing concentrations of ROS and reactive nitrogen intermediates (6).

The intracellular pathogen *Brucella abortus* invades host cells and is trafficked to a compartment known as the Brucella-containing vacuole, where it replicates. To successfully find its way to this niche, *B. abortus* must avoid numerous chemical insults by the host cell and adapt to the nutrient resources available within host cells. The mechanisms by which *B. abortus* evades host defenses and takes advantage of host metabolic responses remain unclear, but previous work showed that *B. abortus* induces a Warburg-like metabolic shift in mammalian host cells. We characterize this host metabolic shift and provide evidence that it is advantageous for *B. abortus* survival in the intracellular niche. We further demonstrate that chemical inhibition of Warburg metabolism attenuates *B. abortus* survival in cultured macrophage-like host cells. Our study informs a model in which *B. abortus* exploits an innate immune metabolic response to support infection.

## RESULTS

### *Brucella abortus* infection of human macrophage-like cells disrupts mitochondrial function and localization

To characterize the effect of intracellular colonization by *B. abortus* on human cell metabolism, we employed mammalian cell Phenotype MicroArrays to measure respiration in terminally differentiated, metabolically active THP-1 cells. These cells were cultivated in minimal essential medium in the presence of central metabolic substrates specifically selected to measure mitochondrial function. Three of the substrates, α-ketoglutaric acid, β-hydroxybutyric acid, and pyruvic acid are metabolized in mitochondria. We assessed the respiration of cultured human cells utilizing only the provided substrate as a primary energy source by using a colorimetric assay that measures the reduction of a redox indicator dye. This assay allowed us to assess the cells’ ability to metabolize substrates across different molecular pathways, and thus provided a measurement of metabolic pathway function upon infection (Figure 1A). We infected the cells with *B. abortus* at increasing multiplicities of infection (MOI). To eliminate the possibility that *Brucella*-induced cell death caused the changes in the metabolic activity of the human cells, we removed medium containing mitochondrial substrates, replaced it with regular growth medium containing glucose, and assessed cell viability using an MTT (3-(4,5-dimethylthiazol-2-yl)-2,5-diphenyltetrazolium bromide) dye-reduction assay. The number of viable cells was statistically equivalent at different *Brucella* MOI (Figure 1B). Uninfected cells utilized all of the mitochondrial substrates (Figure 1C), but the ability to utilize these substrates decreased in cells infected with *B. abortus* (Figure 1D). This decrease was dependent on MOI (Figures 1A). Both infected and uninfected THP-1 cells were incubated in equivalent media containing gentamycin, and thus we attribute the observed metabolic effect to *B. abortus* infection, and not a component of the culture medium. *B. abortus*-infected cells continued to utilize glucose, in contrast to the other substrates (Figure 1C–D). In fact, glucose consumption by infected THP-1 cells increased with increasing MOI (Fig. 1E and 1F).

**FIG 1.**
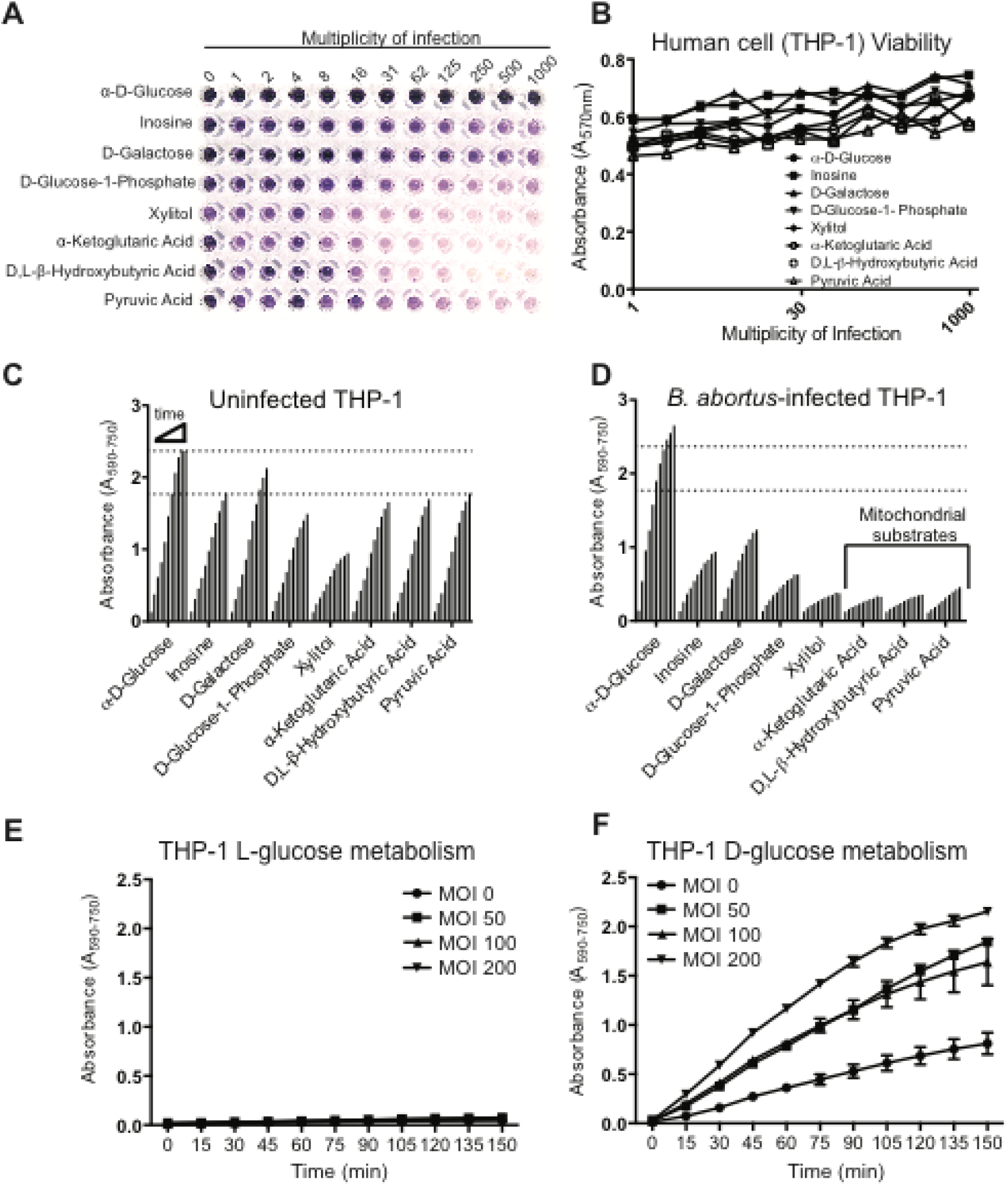
Mitochondrial function is perturbed in THP-1 cells infected with *B. abortus.* **(A)** Mammalian Phenotype MicroArray plate designed to measure mitochondrial function. The ability of THP-1 cells infected with *B. abortus* at increasing multiplicity of infection (MOI) to utilize metabolic substrates was assessed by measuring the reduction of a tetrazolium dye (see Materials and Methods). **(B)** An MTT cell viability assay was performed in the presence of glucose to assess the effect of *B. abortus* infection on human cell survival at the conclusion of the mammalian Phenotype MicroArray assay. Dye reduction by **(C)** uninfected and **(D)** *Brucella*-infected human cells (MOI 125) measured as a time-dependent absorbance readout. Cells were cultured in minimal medium supplemented either with glucose (control) or central metabolic substrates. α-ketoglutaric acid, β-hydroxybutyric acid, and pyruvic acid) that enter the mitochondria, thereby providing a readout of mitochondrial function. Bars along the X axes represent time (0–150 min) in 15-min increments. Dotted lines mark the highest reading for utilization of glucose and pyruvic acid in uninfected THP-1 cells. (E) Metabolic activity of THP-1 in the presence of L-glucose at increasing *B. abortus* MOI monitored by colorimetric dye reduction assay. (F) Metabolic activity of THP-1 in the presence of D-glucose at increasing *B. abortus* MOI monitored by colorimetric dye reduction assay. Panels E and F (*n* = 3); error bars represent standard deviation.

The marked inhibition of mitochondrial substrate utilization suggests that intracellular Brucellae disrupt mitochondrial function. Parasites (7), viruses (8), and intracellular bacterial pathogens including *B. melitensis* (9) and *Listeria monocytogenes* (10) perturb mitochondrial function in host cells. To obtain additional evidence that mitochondrial function is disrupted during *B. abortus* infection, we stained uninfected and infected cells with a dye specific for mitochondria. The flatter morphology of HeLa cells (compared with THP-1) permitted clearer visualization of mitochondria on our microscopy setup in BSL-3 containment, so we used HeLa cells for this assay. Disruptions of overall cell morphology and mitochondrial localization were observed in HeLa cells with detectable *B. abortus*, whereas uninfected cells in the same field of view had normal cell morphology and mitochondrial staining patterns (Figure 2). We do not attribute these changes in mitochondrial staining patterns to gentamicin in the culture medium, as mitochondria were indistinguishable from a control culture lacking gentamicin at concentrations up to 200 µg mL^−1^ (Figure S1). Though infected cells had a more rounded morphology in some cases, an MTT cell viability assay revealed no loss of host cell viability with increasing *B. abortus* MOI within the same timeframe (Figure 1B). Therefore, it is unlikely that the changes in cell morphology and mitochondrial localization observed by fluorescence microscopy are a result of host cell death upon infection.

**FIG 2.**
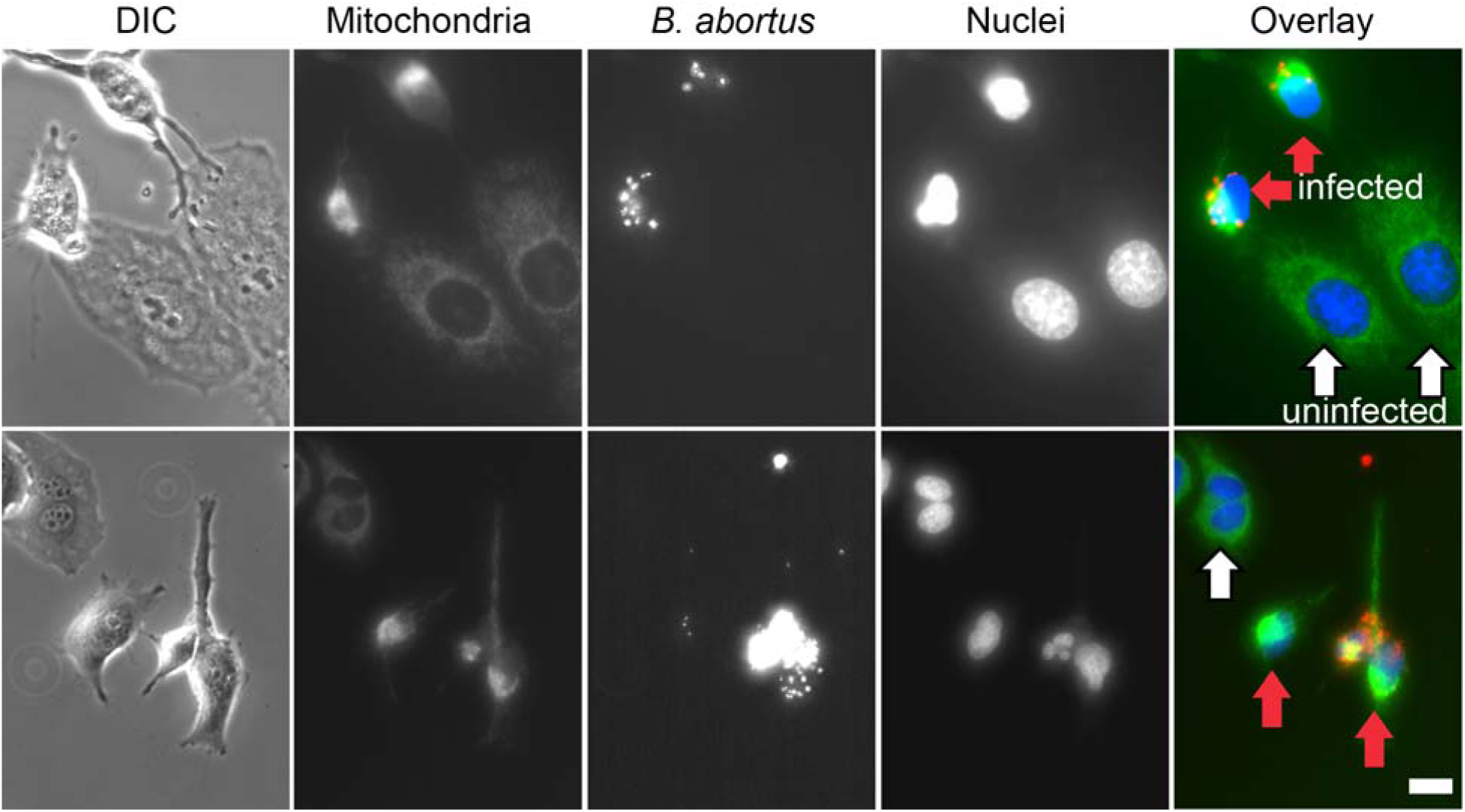
Intracellular *B. abortus* induces morphological changes of mitochondria localization in HeLa cells. Two fields showing mitochondrial staining (MitoTracker, green), *B. abortus*-Tn7mCherry (mCherry fluorescence, red), nuclei (Hoechst stain, blue), and the corresponding image with differential interference contrast (DIC) microscopy. Red arrows indicate cells with visible intracellular *B. abortus*, white arrows indicate cells for which no infection is evident. Cells not infected with *B. abortus* have typical mitochondrial staining patterns (also see Figure S1). Scale bar = 50 µm.

Most amino acids can be metabolized to enter the TCA cycle through anaplerotic reactions (11). To determine whether the observed *B. abortus*-mediated disruption of host mitochondrial function directly affects amino acid metabolism, we assessed the effect of infection on the host’s ability to utilize amino acids as a carbon source. While glucose metabolism was unaffected, infected cells could not metabolize any of the amino acids known to enter the TCA cycle (Figure 3). Together, the results of these experiments are consistent with a model in which infection of human cells with *B. abortus* results in major metabolic changes that are linked to perturbation of mitochondrial function.

**FIG 3.**
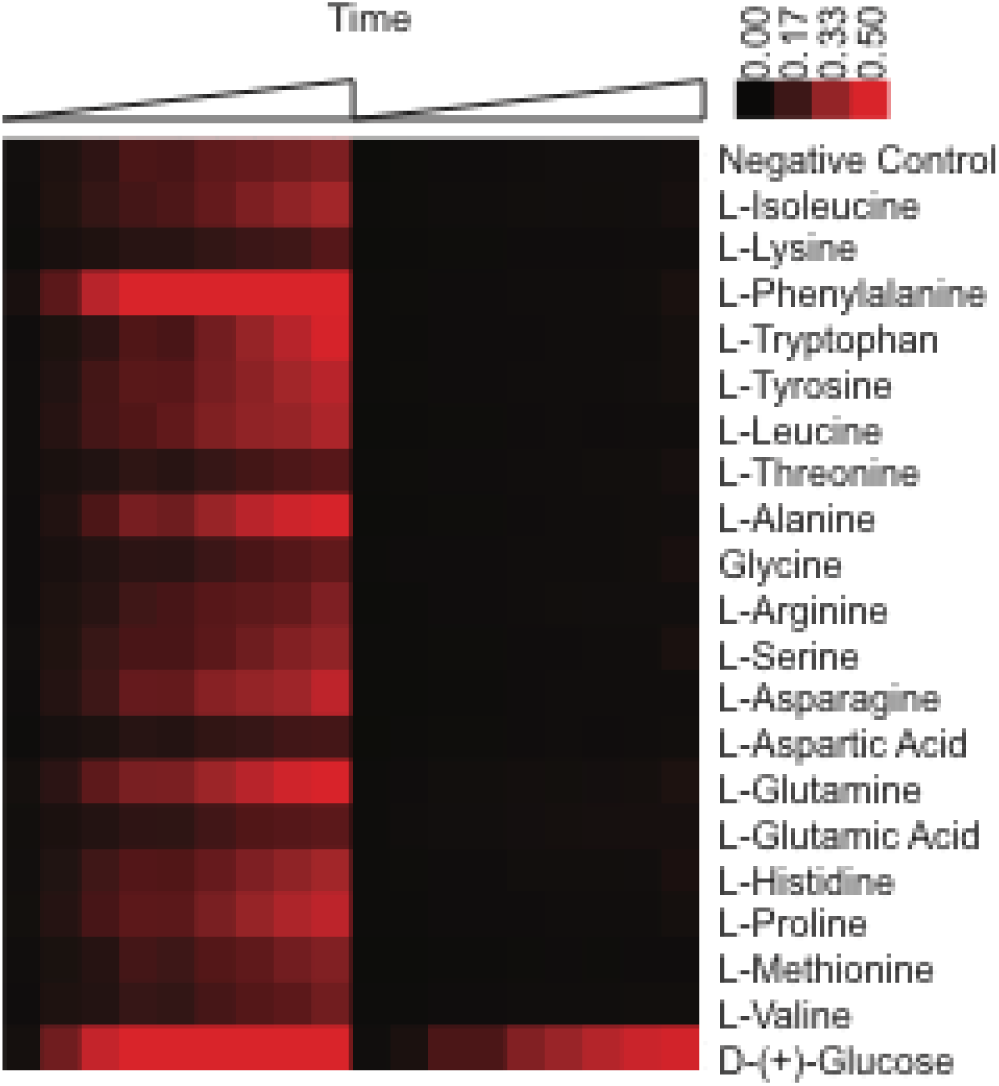
Host cell utilization of amino acids as a carbon source before and after *B. abortus* infection. Heat map representing host cell metabolism as measured colorimetrically via a redox dye (see Materials and Methods). Time-dependent metabolic change of non-infected versus *B. abortus*-infected THP-1 cells in the presence of amino acids are plotted. Color is an arbitrary scale from black (no reduction of the dye, 0.00) to red (maximum reduction of the dye, 0.50). THP-1 has low metabolic activity in the PM panel negative control, which is presumed to be due to utilization of component(s) of fetal bovine serum. Addition of *B. abortus* inhibits THP-1 metabolism in this condition and in all assayed amino acids, but not in glucose (see Figure 1F for *B. abortus* MOI dependence of host glucose metabolism).

### *Brucella*-dependent induction of lactic acid production in human cells

Macrophages and dendritic cells have central functions in innate immunity. Upon activation, they can undergo a shift in energy metabolism by shutting down oxidative phosphorylation and increasing the rate of aerobic glycolysis in a process known as the Warburg effect (3). One hallmark of this effect is the production of lactic acid in the presence of oxygen. A previous study noted an increase in lactate production in non-polarized macrophages infected with *Brucella* spp. (12). To test whether THP-1 infected with *B. abortus* shift their metabolism toward a Warburg-like state, we measured the production of lactic acid by infected cells and found that in THP-1 cells, lactic acid production increased with increasing MOI (Figure 4A). To test whether this metabolic shift required infection with live *B. abortus*, we measured lactate production by THP-1 cells exposed to heat-killed *B. abortus*. Heat-killed *Brucella* induced similar lactate production from the THP-1 human cell line, although at the highest MOI, the THP-1 cells treated with heat-killed pathogen produced less lactate, compared with cells infected with live pathogen (Figure 4B). These results, together with the data presented in Figures 1–3 support a model in which a molecular component of the *B. abortus* cell promotes a Warburg-like shift in host cell metabolism by disrupting mitochondrial function.

**FIG 4.**
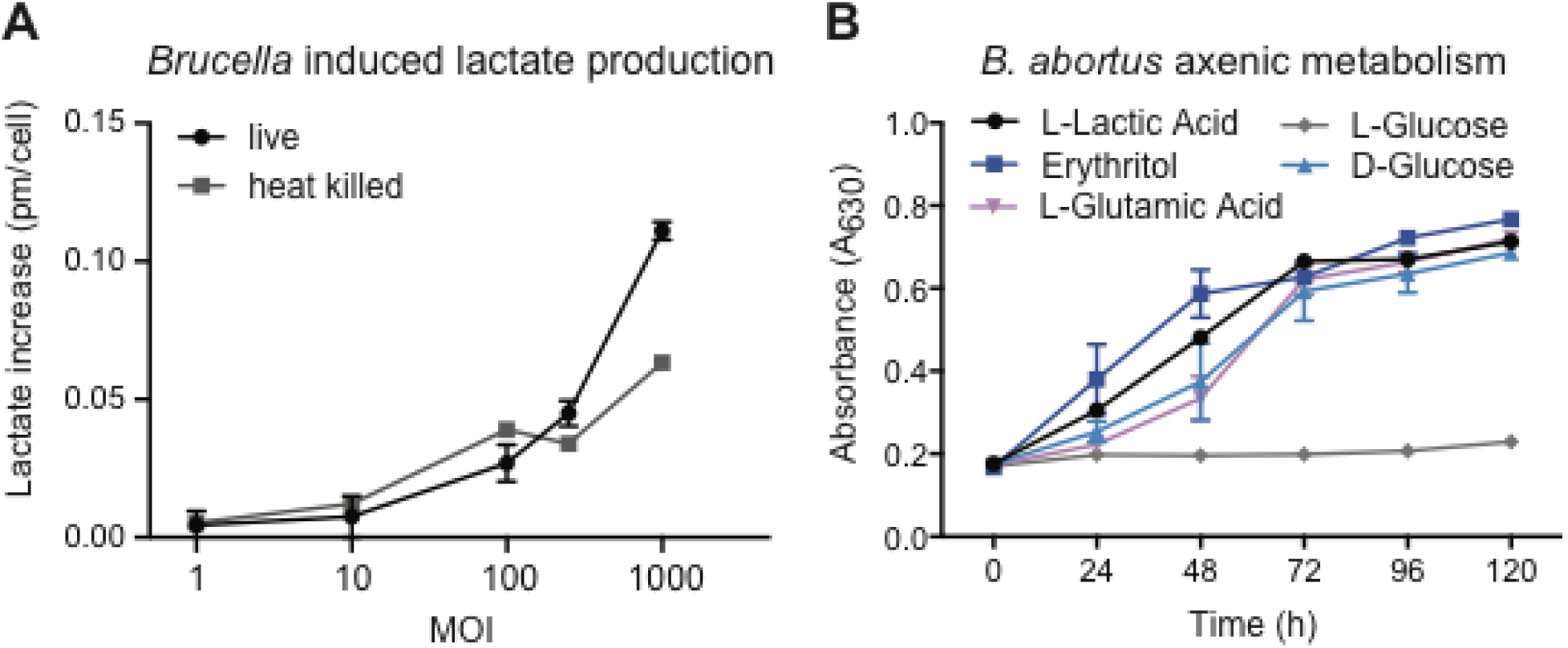
*B. abortus* induces host cell lactate production and utilizes lactate as a carbon and energy source as efficiently as glucose, erythritol, and glutamate. **(A)** Lactate levels in THP-1 cells treated with live or heat-killed *B. abortus* at different multiplicities of infection. **(B)** Metabolism of *B. abortus* measured on carbon/energy sources L-lactate, L-glutamate, erythritol, D-glucose, and L-glucose. Data show an average of two independent experiments ± standard deviation.

### *Brucella abortus* requires lactate dehydrogenase for normal intracellular survival in THP-1 cells

*Brucella* and *Salmonella* both replicate intracellularly and can persist in the host for long periods by shifting host metabolism to increase the available glucose (12–14). We measured *B. abortus* growth in axenic culture on lactic acid, compared with three substrates that it can efficiently metabolize, L-glutamic acid, glucose, and erythritol. We found that *B. abortus* grows on lactic acid as the sole carbon and energy source as well as it does on glutamic acid, glucose, or erythritol (Figure 4B). We postulated that, during infection, *B. abortus* may exploit the Warburg-type shift in host metabolism by utilizing lactic acid as a carbon and energy source.

To further assay *B. abortus* lactic acid metabolism, we engineered a strain harboring an in-frame deletion of the gene encoding L-lactate dehydrogenase (LDH), *Bab2_0315* (Δ*lldD*). To determine whether deletion of *lldD* affects utilization of other carbon sources, we used Phenotype MicroArrays to compare the metabolic activity of Δ*lldD* to wild type, and found that utilization of L-lactic acid was the most statistically significant loss-of-function phenotype (Figure 5A). The wild-type *B. abortus* exhibited robust growth on lactic acid as a carbon and energy source, but Δ*lldD B. abortus* did not grow on lactic acid (Figure 5B). The Δ*lldD* mutant had no apparent growth defect in *Brucella* broth supplemented with glucose (Figure 5C), but the numbers of colony forming units (CFU) of Δ*lldD* were significantly reduced in THP-1 macrophages relative to wild type at the 48-hour time point (Figure 5D). Complementation of the mutation by introduction of *lldD* at its native locus rescued the THP-1 replication/survival defect. Together, our results indicate that, upon infection, human cells undergo a metabolic shift that results in increased lactate production. The lactate catabolic enzyme LDH is required for normal intracellular *B. abortus* replication and/or survival in an *in vitro* macrophage infection model.

**FIG 5.**
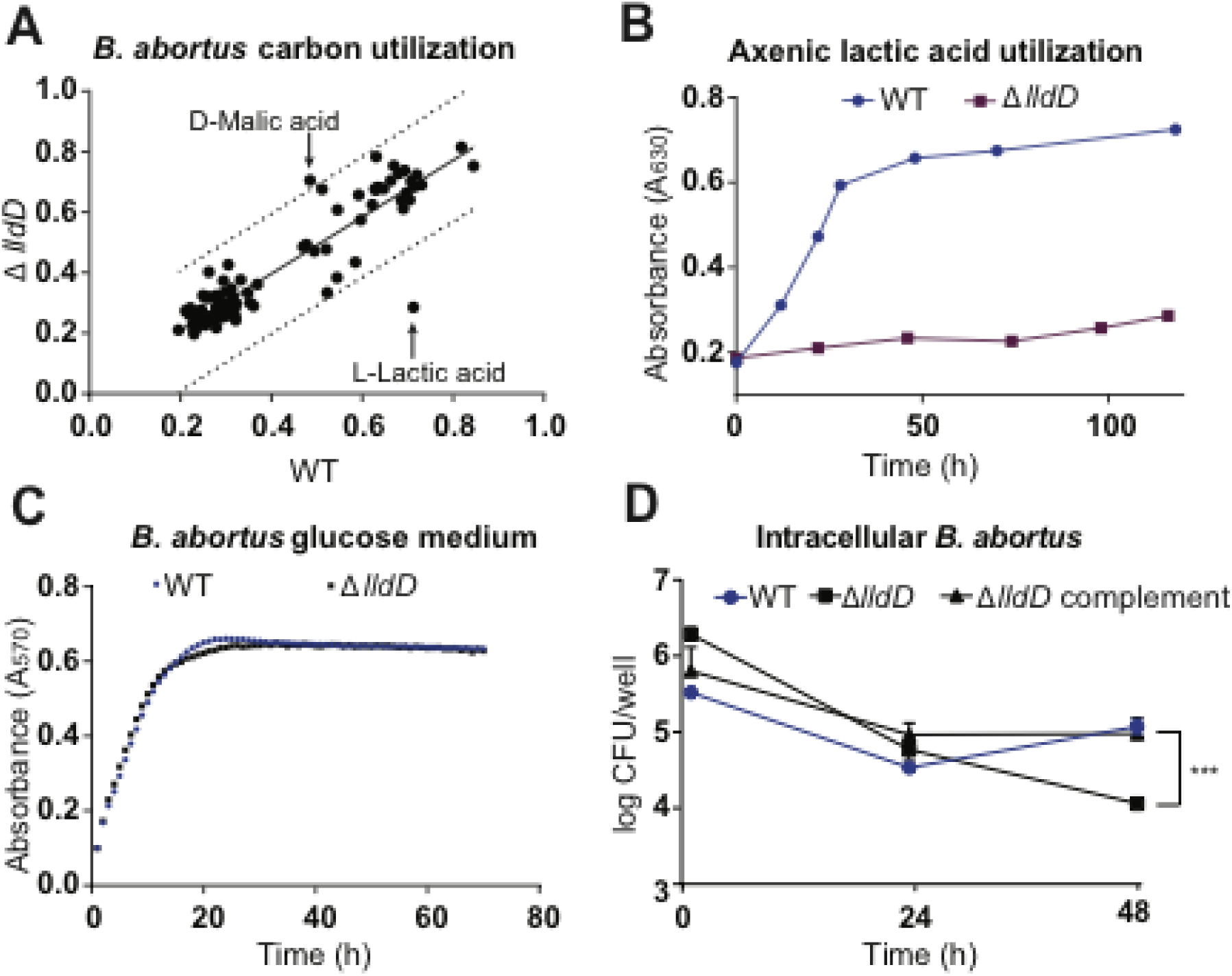
*B. abortus* L-lactate dehydrogenase deletion strain Δ*lldD* cannot utilize lactic acid as a sole carbon and energy source in axenic culture and is attenuated in a macrophage infection model. **(A)** Phenotype MicroArray carbon utilization measurements of wild-type (WT) *B. abortus* and L-lactate dehydrogenase (LDH) mutant (Δ*lldD*) strains. Metabolic activities for growth on 95 distinct carbon substrates (see Materials and Methods) are plotted against each other; the dotted line marks a 99% prediction band of a linear regression. Correlation values above the line represent a gain-of-function phenotype and values below the line represent a loss-of-function phenotype. L-lactic acid and D-malic acid are marked as loss-of-function and gain-of-function, respectively **(B)** Graph representing time-dependent utilization of lactic acid as a carbon energy source, and measured by a metabolic indicator dye (see Materials and Methods). **(C)** Graph representing axenic growth of the WT and Δ*lldD* strains in glucose-containing *Brucella* broth. **(D)** Intracellular replication and survival of the WT and Δ*lldD* strains in THP-1 macrophage cell infection model. Intracellular *B. abortus* were quantified as log_10_ colony forming units (CFU) per well at 1, 24, and 48 h post-infection. Lines represent WT (•), Δ*lldD* mutant (▪), and complement (▲). *n* = 3, Error bars represent standard deviation.

### Small-molecule glycolysis inhibitors decrease intracellular replication of *Brucella abortus* but do not affect axenic growth

Identification of pathogen-induced changes in host metabolism offers new avenues for the development of anti-infectives. Since *Brucella*-infected cells have disrupted mitochondrial metabolism and increased glycolysis, we used small-molecule inhibitors of glucose metabolism (Figure 6A) to determine whether inhibition of the host glycolytic pathway and lactate production affected intracellular replication/survival of *B. abortus*. We tested: 3-bromopyruvic acid (3-BPA), a potent glycolytic inhibitor (15, 16); methyl 1-hydroxy-6-phenyl-4-(trifluoromethyl)-1H-indole-2-carboxylate (NHI-2), an inhibitor of lactate dehydrogenase A (LDH-A) (17); and 2-deoxy-D-glucose (2-dG), a non-metabolizable analog of D-glucose (18). We counted *B. abortus* cells located inside THP-1 cells treated with each of these anti-metabolic compounds at various concentrations and expressed the results as the number of *Brucella* per THP-1 nucleus. Fluorescent *Brucella* puncta were often larger than a single *B. abortus* cell, particularly in untreated cells, and likely contained multiple bacteria. Nonetheless, we took a conservative approach and counted each fluorescent *B. abortus* spot as one cell. Each anti-metabolic drug significantly inhibited replication/survival of intracellular *Brucella* in live cells (Figure 6B). This effect was particularly evident when drug-treated and untreated cells were visualized by fluorescence microscopy (Figure 6E).

**FIG 6.**
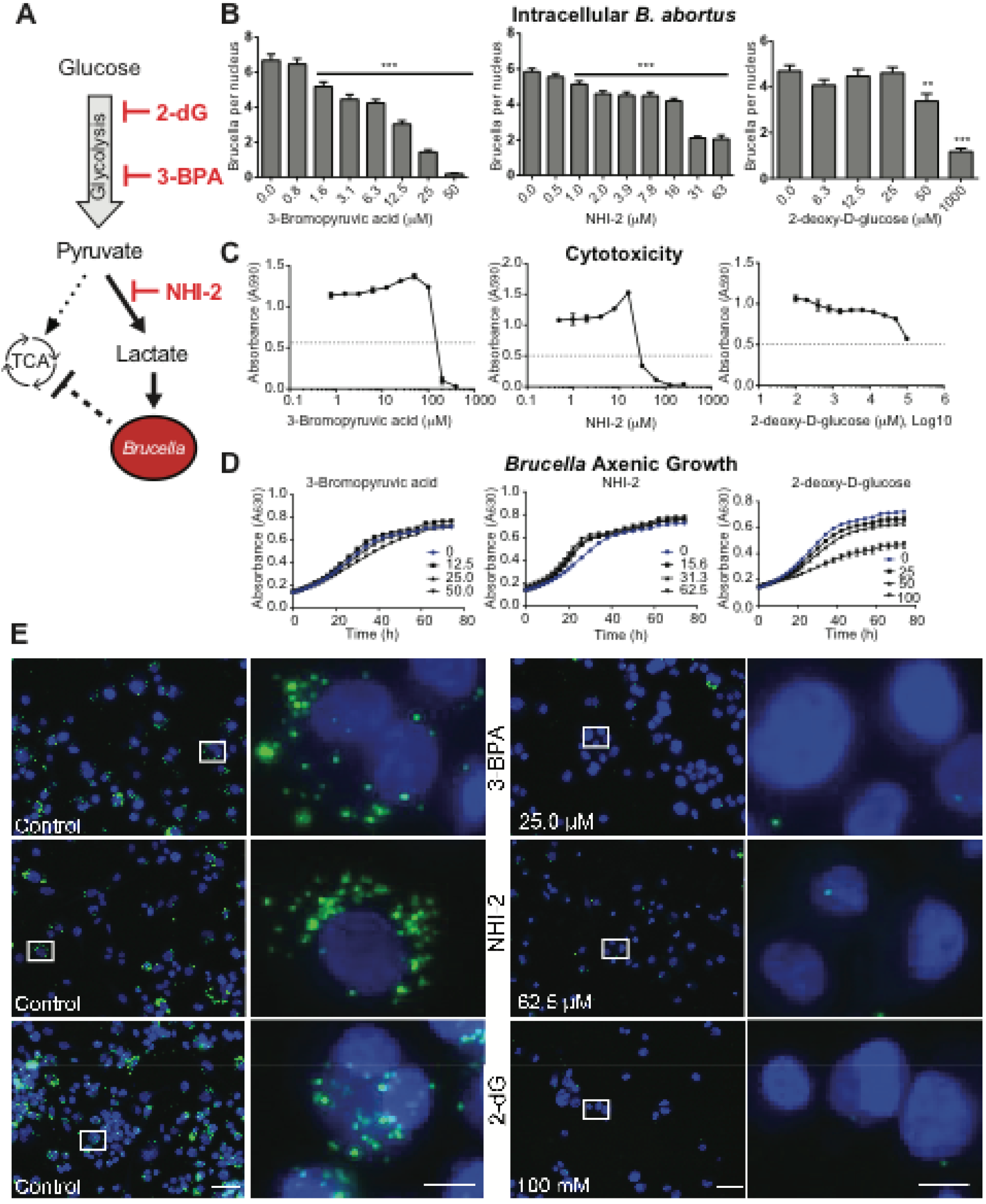
Pharmacological disruption of the host glycolytic pathway and lactate production inhibits replication of intracellular *B. abortus*. **(A)** Model depicting *Brucella*-dependent modulation of carbon metabolism and activation of aerobic glycolysis, resulting in the production of lactate. *B. abortus* utilizes lactate as efficiently as preferred substrates glucose and erythritol. Pharmacological inhibitors of glycolysis, 3-Bromopyruvic acid (3-BPA), NHI-2, and 2-deoxy-D-glucose (2-dG), inhibit intracellular growth of *B. abortus*: **(B)** Graphs represent the effect of 3-BPA, NHI-2, and 2-dG on intracellular replication of *B. abortus* as quantified by enumerating fluorescent bacteria per THP-1 nucleus; the effect of anti-glycolytic compounds on infection was assessed by measuring the average ratio of bacteria-to-nuclei in THP-1 cells treated with each of the compounds. Statistical significance was evaluated from four biological replicates (10 images each) using one-way analysis of variance (ANOVA) followed by Dunnett’s post test \*\**p* < 0.01, \*\*\**p* < 0.001). Error bars represent ± standard deviation from the mean. **(C)** Cytotoxicity of inhibitory compounds to THP-1 cells; an XTT assay was used to measure compound cytotoxicity to THP-1 cells. **(D)** Effect of metabolic inhibitors on *B. abortus* axenic growth/metabolism in axenic culture. *Brucella* growth/metabolism was assayed by measuring tetrazolium-based dye reduction at630 nm **(E)** Fluorescence images of nuclei (blue) and intracellular *B. abortus* (pseudocolored green) reflect the efficacy of 3-BPA, NHI-2, and 2-dG to inhibit *B. abortus* replication in the intracellular niche. Scale bars = 10 µm (full view) and 50 µm (magnified insets, marked with white box in full view). Dotted line = half maximal inhibitory concentration (IC50).

To determine whether the antimicrobial activity of these compounds coincides with their cytotoxicity to human cells, we assessed their cytotoxicity using an XTT (2,3-Bis-(2-Methoxy-4-Nitro-5-Sulfophenyl)-2H-Tetrazolium-5-Carboxanilide) cell viability assay. As expected, each metabolic inhibitor was cytotoxic at high concentrations (Figure 6C). However, treatment of THP-1 cells with 3-BPA, the most potent compound, inhibited *B. abortus* intracellular replication/survival at concentrations that were well below the THP-1 cytotoxicity threshold. NHI-2 and 2-dG were less potent, but effectively inhibited replication/survival of intracellular *Brucella* (Figure 6B and 6C). Both 3-BPA and NHI-2 significantly inhibited intracellular *Brucella* at concentrations below the cytotoxicity threshold.

Finally, to determine whether the antimicrobial effect of these compounds occurs by targeting host metabolism, or whether they also directly inhibit bacterial growth, we measured growth and viability of *B. abortus* in axenic culture in the presence of 3-BPA, NHI-2, and 2-dG. Neither 3-BPA nor NHI-2 affected growth and viability of *Brucella* in axenic medium, though 2-dG had an inhibitory effect on axenic *B. abortus* growth at higher concentrations (≥ 100 mM).

These data support a model in which inhibition of host glycolysis and lactate production with 3-BPA and NHI-2 inhibits intracellular *B. abortus* replication/survival without directly targeting *B. abortus* growth (Figure 6D). Together, these results demonstrate that infection-induced changes in host central metabolic pathways are required for proper replication/survival of intracellular *B. abortus*. Inhibition of the glycolytic pathway with host-targeting anti-metabolic drugs may provide an effective means to decrease intracellular bacterial load without directly targeting the bacterium (Figure 6E).

## DISCUSSION

We have identified large metabolic changes in THP-1 host cells that occur upon exposure to the intracellular bacterial pathogen *B. abortus*. Intracellular *B. abortus* clearly perturb the function of host mitochondria, as characterized by re-localized mitochondrial staining and the loss of the ability to utilize mitochondrial substrates (Figure 1 and 2). The host metabolic effects we observed upon *B. abortus* infection resemble the effect of mitochondrial inhibitors (19), and are consistent with the previously reported down-regulation of mitochondrion-associated gene expression in macrophages infected with *B. melitensis* (20). Many bacteria are known to affect host mitochondria (21), but only a few species have been reported to disrupt mitochondrial structure or localization (9, 10, 22). The mitochondrial staining profile of infected cells observed in our study resemble those of cells treated with uncouplers in combination with a respiration inhibitor; although this treatment results in energy deprivation and aggregation of mitochondria, cells remained viable in this assay (23).

Intracellular pathogens compete with host cells for available nutrients, including central metabolites and amino acids (24). Human THP-1 cells utilize amino acids as an energy source, but this is inhibited by *B. abortus* infection (Figure 3). The reduced ability of infected host cells to utilize amino acids suggests a model whereby intracellular *B. abortus* competes for these substrates early in infection by diverting the host’s metabolism away from amino acid consumption. This model is consistent with proteomic data showing down-regulation of carbohydrate catabolic proteins and enhanced expression of enzymes involved in amino acid catabolism several hours after macrophage infection (25). Our data are also congruent with a recent model from Zúñiga-Ripa and colleagues (26) based on genetic analyses of *B. abortus* metabolic mutants. Data from these investigators suggest that pentose and hexose sugars are limited in the host and that energy is compensated by amino acids in certain infection niches.

In addition to our investigation of shifts in host amino acid metabolism upon infection, we also explored the induction of host lactate production upon exposure to *B. abortus*. Erythritol, glutamic acid, and glucose are efficiently metabolized by *Brucella* (27, 28), and our results show that, in axenic culture, *B. abortus* strain 2308 can metabolize lactate as efficiently as these substrates as the sole carbon and energy source (Figure 4). Our data are thus consistent with a study from Gerhardt and Wilson reporting that lactate can be utilized as a carbon and energy source by *Brucella abortus* strain 19 (29), though the level of growth we observed for strain 2308 in the defined Phenotype MicroArray-lactate medium was significantly higher than that observed in this early report. *Brucella* spp. may take advantage of host lactate as an energy source in niches where lactate is naturally abundant such as the placenta (30) or male reproductive tissue (31). Both lactate and amino acids have been proposed to provide energy to certain intracellular pathogens within the host environment (32). *Brucella* spp. pathogens have a tropism for genital organs, and the possibility that lactate may serve as an important metabolic substrate for *Brucella* spp. during colonization of the reproductive tract has been recently reviewed by Letesson and colleagues (33).

Our observations that *B. abortus (a)* efficiently metabolizes lactate as a carbon and energy source and *(b)* that THP-1 host cells exposed to *B. abortus* increase glucose consumption and lactate production raises the possibility that *Brucella* spp. takes direct advantage of a Warburg-like shift in host inflammatory cells to support growth during infection. We took two approaches to test whether host lactate affects intracellular growth and/or survival in THP-1 macrophages. First, we created a *B. abortus* lactate dehydrogenase knockout strain Δ*lldD*, which cannot utilize lactate in Δ axenic culture (Figure 5A-B). The Δ*lldD* mutant showed no growth defects across a range of axenic media, but its intracellular replication/survival was attenuated by ≈ 1 log after 48 h in host THP-1 cells (Figure 5D). This provides evidence that the ability to utilize lactate is necessary for survival in the intracellular niche in the THP-1 model. In a second approach, we targeted host lactate synthesis using the small molecules NHI-2, 3-BPA, and 2-dG, which decrease lactate levels and inhibit aerobic glycolysis (17, 34, 35). Each of these metabolic inhibitors decreased intracellular growth and/or replication of *Brucella* (Figure 6). These compounds are known for their anticancer properties, but their anti-infective properties were not previously reported.

The Warburg shift in host metabolism upon pathogen infection is a well-documented effect. For example, *Mycobacterium tuberculosis* enhances glycolysis in THP-1 cells (36). Early studies of infection-induced changes in host metabolism showed that cells infected with *Chlamydia* increase utilization of glucose through glycolysis and increase lactate production (37). Other pathogens, including the parasite *Plasmodium falciparum* (38), and Mayaro virus (39), induce similar changes in host metabolism. Given these results, it is unlikely that the changes observed in our study are specifically induced by *B. abortus*. This is supported by our observation that lactate production is induced upon exposure to heat-killed *B. abortus* (Figure 4A).

The relevance of a *Brucella*-induced increase in aerobic glycolysis and lactate production to *in vivo* infection biology in natural hosts remains unclear. The importance of *B. abortus* glycolysis in infection is supported by data showing that deletion of pyruvate kinase, which presumably renders the strain unable to execute the last step of glycolysis, results in attenuation in macrophage and mouse infection models (40). However, this mutation is pleiotropic and may result in attenuation for a variety of reasons. While classically activated M1 macrophages undergo a shift to Warburg-like metabolism upon exposure to pathogen-associated molecular patterns (PAMPs), Xavier and colleagues have shown that *B. abortus* preferentially infects alternatively activated M2 macrophages in mice, which have an opposite metabolic profile with decreased aerobic glycolysis (12). As recently noted (26, 33), the physiological characteristics of host cell types and tissues colonized by *Brucella* spp. during the course of disease vary significantly, and it may be that lactate catabolism is important in some *in vivo* microenvironments but irrelevant in others.

Collectively, the data presented in this study support a model in which intracellular *B. abortus* can subvert host amino acid and carbohydrate metabolic pathways to support its growth and survival in the host niche (Figure 7). The host anti-metabolic compounds identified here and in previous studies (41) may provide a novel therapeutic approach to treat *B. abortus* and other bacterial infections that rely on host metabolic intermediates.

**FIG 7.**
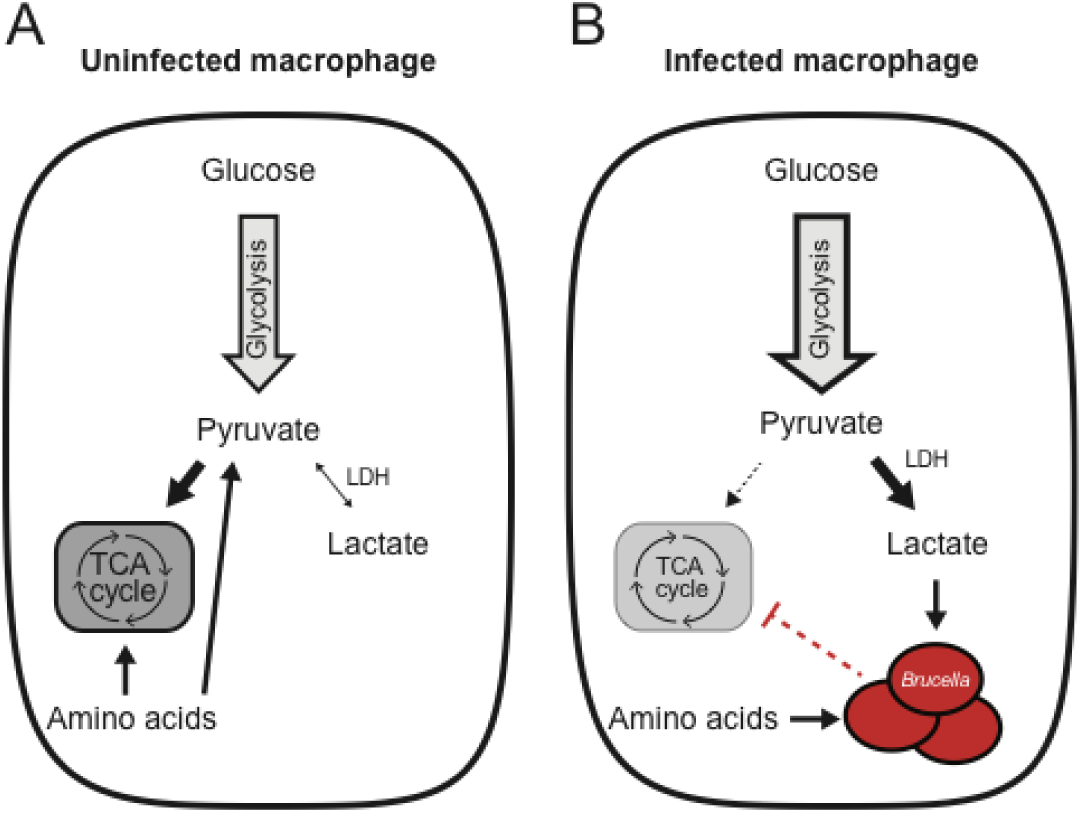
Model of how *B. abortus* may exploit a shift to Warburg-like metabolism in host cells. Data presented in this study support a model in which inhibition of tricarboxylic acid (TCA) cycle metabolism upon infection biases the host cell away from amino acid catabolism and toward the production of lactic acid. **(A)** Uninfected macrophages exhibit typical central carbon metabolism in which pyruvate enters the TCA cycle. **(B)** *B. abortus* infection biases the macrophage toward aerobic glycolysis and lactate production and decreased amino acid catabolism. *B. abortus* can utilize select amino acids as carbon and energy substrates and can also utilize lactic acid via the lactate dehydrogenase (LDH) enzyme, LldD. A strain lacking *lldD* is compromised in lactic acid metabolism in axenic culture and in intracellular survival in a THP-1 macrophage model. Pharmacological inhibition of host glycolysis or LDH-A reduces *B. abortus* survival in the intracellular niche in a THP-1 infection model.

## MATERIALS AND METHODS

### Bacterial strains

All studies on live *B. abortus* strain 2308 were conducted at Biosafety Level 3 (BSL3) at the Howard Taylor Ricketts Regional Biocontainment Laboratory, University of Chicago, according to US Federal Select Agent Program guidelines. *B. abortus* expressing mCherry was previously generated from the wild-type *B. abortus* 2308 parent strain by integrating mini-Tn7 expressing a single copy of *mCherry* at the *glmS* locus. Bacterial strains used in this study are listed in Table S1.

### Mammalian tissue culture

Prior to each experiment, THP-1 macrophage-like cells were grown to a maximum density of 1 × 10^6^ mL^−1^ in complete RPMI-1640 medium (HyClone) supplemented with 2 mM glutamine (Gibco) and 10% heat-inactivated fetal bovine serum (FBS) (HyClone). HeLa cells were grown in Dulbecco Modified Eagles Medium (DMEM; HyClone) supplemented with 10% heat-inactivated FBS. All tissue culture cells were grown at 37°C in a humidified environment with 5% CO_2_.

### Mammalian Phenotype MicroArray: mitochondria toxicity

THP-1 cells were seeded at 50,000 cells/well in two half-area 96-well plates in the presence of 40 ng mL^−1^ phorbol myristate acetate (PMA), and allowed to differentiate for 72 h at 37°C. One plate was infected, while the other was a non-infected control. *B. abortus* 2308 strain was streaked onto Schaedler blood agar (SBA) plates before growing for 2 d, re-streaking, and growing for another 2 d. On the day of infection, *Brucella* cells were scraped off the SBA plate, and suspended and washed in RPMI-1640 growth medium. Cell concentration was measured by the optical density method at 600 nm (OD_600_). For the infection plate, bacteria were added to the last column of the 96-well plate at 1000 MOI, and serially diluted by a factor of two. The first column was left uninfected. Both uninfected and infected plates were centrifuged at 2170 rpm for 10 min, followed by a 1-h incubation at 37°C and 5% CO_2_. Gentamicin was added to each well of both uninfected and infected plates at a final concentration of 100 µg mL^−1^ and the plates were further incubated for 1 h. Following incubation, the medium was removed from both plates and replaced with fresh medium containing 25 µg mL^−1^ gentamicin. Cells were incubated for 24 h. Twenty-four hours after infection, two PM-M TOX1 MicroPlates (Biolog) were reconstituted with 50 µL (per well) of MC-0 medium (Biolog IF-M1 medium, 0.3 mM L-Gln, 5% dialyzed FBS, 25 µg mL^−1^ gentamicin). The culture medium was removed from infected cells and replaced with the contents of reconstituted PM-M TOX1 plates. *Brucella*-infected cells were incubated in the presence of mitochondrial substrates for another 24 h. Following this incubation, 10 µL of PM-A dye (Biolog) was added to each well, and the absorbance measured every 15 min (up to 150 min) at 590 nm and 750 nm using a Tecan Infinite 200 PRO microplate reader. Plates were incubated at 37°C and 5% CO_2_ between readings. The collected absorbance 590 nm data (minus reference absorbance at 750 nm) were analyzed as a function of time.

Following the experiment, cell viability was assessed by removing all liquid from the PM-M TOX1 plate, replacing it with glucose-supplemented growth medium containing 10% FBS, and 0.5 mg mL^−1^ MTT reagent, then measuring the absorbance at 570 nm after 24 h.

### Mammalian Phenotype MicroArray: utilization of amino acids

THP-1 and *B. abortus* 2308 cells were prepared as described above. The infection protocol was as described above with small changes. Bacteria were added to differentiated THP-1 cells in a 96-well plate at 100 MOI. Plates were centrifuged at 2170 rpm for 10 min followed by 1-h incubation at 37°C and 5% CO_2_. Gentamicin was added to each well of uninfected and infected plates at a final concentration of 100 µg mL^−1^ and plates were further incubated for 1 h. Following incubation, the medium was removed from both plates and replaced with fresh medium containing 25 µg mL^−1^ gentamicin. Cells were incubated for 24 h. Twenty-four hours after infection, the content of the mammalian Phenotype MicroArray PM-M2 (Biolog) was reconstituted with 50 µL (per well) of MC-0 medium (Biolog IF-M1 medium, 0.3 mM L-Gln, 5% dialyzed FBS, 25 µg mL^−1^ gentamicin). The culture medium was removed from infected cells and replaced with the contents of reconstituted PM-M2 plates. *Brucella*-infected cells were incubated in the presence of various amino acids for another 24 h. Following this incubation, 10 µL of dye PM-A (Biolog) was added to each well, and the absorbance was measured every 15 min (up to 120 min) at 590 nm and 750 nm using a Tecan Infinite 200 PRO microplate reader. Plates were incubated at 37°C and 5% CO_2_ between readings. Absorbance at 590 nm data (minus reference absorbance at 750 nm) was analyzed as a function of time. Results were analyzed by hierarchical clustering using Gene Cluster 3.0 and visualized using Java TreeView (v1.1.6r4).

### Bacterial Phenotype MicroArray

Utilization of glucose, erythritol, and lactic acid was measured using the Phenotype MicroArray as previously described (42). Briefly, *B. abortus* strains (wild type 2308 and Δ*bab2_0315*) were streaked onto an SBA plate, cultivated at 37°C and 5% CO_2_ for 48 h, re-streaked, and grown for another 48 h. Cells were scraped off the plate and resuspended in 1 x IF-0a medium (Biolog) at a final density equivalent to 5% transmittance (OD_600nm_). Phenotype MicroArray inoculating fluids (PM1 and PM2) were prepared from 0.2-µm-filtered and sterilized stock solutions at the following final concentrations: 2 mM MgCl_2_, 1 mM CaCl_2_, 25 μM l-arginine, 50 μM l-glutamic acid, 5 μM β-NAD, 25 μM hypoxanthine, 25 μM 5′-UMP, 25 μM l-cystine (pH 8.5), 0.005% yeast extract, and 0.005% Tween 40. PM1 and PM2 inoculating fluids were prepared in 1 x IF-0a GN/GP medium (Biolog). Dye mix G (Biolog) was added to each solution to a final concentration of 1X. A 1:13.6 dilution of 5% transmittance *Brucella* suspension was added to each inoculating fluid. Per well, 100 µL of *Brucella*-containing inoculating fluid was dispensed into each plate. Phenotype MicroArray plates were incubated at 37°C and 5% CO_2_, and the absorbance at 630 nm was measured every 24 h for up to 5 d, using a Tecan Infinite 200 PRO microplate reader.

### Imaging of mitochondria

HeLa cells were seeded onto 22 × 22 mm cover slides at 50,000 cells/well in a 6-well plate in 2 mL of DMEM medium per well. Cells were grown for 24 h prior to infection. Prior to infection, a culture of *B. abortus* constitutively expressing mCherry was inoculated in *Brucella* broth, and grown overnight at 37°C on orbital shaker. On the day of infection, *Brucella* cells were collected, resuspended in DMEM, and opsonized by incubating for 30 min at 37°C with 1:1000 dilution of serum obtained from *Brucella*-infected mice, as previously described (43). Following opsonization, cells were washed and added to HeLa cells at 10,000 MOI. The plate was centrifuged at 2170 rpm for 20 min, and placed for 24 h at 37°C and 5% CO_2_. Following incubation, cells were washed with DMEM, re-infected at 10,000 MOI, and incubated for another 24 h. After this incubation, cells were washed with DMEM, the cover slide was transferred to a 50 mL tube containing 15 mL DMEM and 200 µg mL^−1^ gentamicin. The slide was incubated at 37°C and 5% CO_2_ for 1 h. Following the incubation, 15 µL of 10 mg mL^−1^ Hoechst stain and 2 µL of 1 mM MitoTracker (Molecular Probes) dye were added and incubated for another 20 min. The cover slide was washed and placed on a microscope slide with cells facing up. A drop of DMEM was placed on top of the cover slide and covered by another 22 × 22 mm cover slide. To prevent medium evaporation, the edges were quickly sealed with nail polish, and live cells were immediately imaged using a Leica-DMI6000B fluorescence microscope equipped with a HC PL APO 63X/1.4 NA Oil PH3 CS2 objective. Images were captured using Leica Application Suite X and Hamamatsu Orca-R2 camera. To test the effect of gentamicin on mitochondria staining, HeLa cells were treated with 0 to 200 µg mL^−1^ gentamicin for 24 h and stained as described above.

### Quantification of lactic acid

Intracellular lactic acid levels were quantified according to the manufacturer’s protocol using a lactate assay kit (Sigma-Aldrich). Briefly, THP-1 cells were seeded and infected with either live or heat-killed *B. abortus* at increasing MOI, as described above. Infected cells were incubated for 48 h before quantifying lactate levels. After incubation, cells were washed with PBS, homogenized in 60 µL of lactate assay buffer, and cell debris was removed by centrifuging at 14,000 *g* for 10 min. Fifty microliters of supernatant was collected from each sample, and endogenous lactate dehydrogenase was removed by centrifuging supernatants at 14,000 *g* for 30 min using 10 kDa MWCO Ultra-0.5 mL spin filter (Amicon). Twenty microliters of lactate assay buffer was added to each sample to bring the volume up to 50 µL. Fifty microliters of master mix (lactate assay buffer, enzyme mix, and probe) was added to each well, mixed, and incubated in the dark at room temperature for 30 min before measuring absorbance at 570 nm. Lactate concentration was calculated based on a standard lactate curve and normalized to the number of cells per well. To assess the effect of heat-killed *B. abortus* on THP-1 lactate production, *B. abortus* cells were first titered by serial dilution. These titered cells were then incubated for 1 hour at 95 degrees C to heat kill.

### Construction of *B. abortus lldD* chromosomal deletion

A chromosomal deletion of *B. abortus lldD* (*bab2_0315*) was obtained using a double recombination strategy. Primers used in this study are given in Table S2. The 500 base pairs upstream (using primers 1775 and 1779) and downstream (using primers 1776 and 1777) of *lldD* were Gibson-cloned into pNPTS138, which was digested with *Bam*HI and *Hin*dIII (New England Biolabs). Constructs were validated by DNA sequencing. To delete *lldD*, the deletion plasmid was transformed into *B. abortus*, and primary integrants were selected by plating on SBA supplemented with 50 µg mL^−1^ kanamycin (SBA-Kan). The resulting colonies were grown overnight in *Brucella* broth without selection and plated on SBA supplemented with 5% sucrose to select for second crossovers. The resulting colonies were screened for the *lldD* deletion by sequencing using the sequencing primers 1788/1789.

To complement the *lldD* deletion strain, the *lldD* gene plus the 500 bp upstream and downstream (using primers 1775/1777) were Gibson-cloned into pNPTS138, and restored at the native site using a two-step recombination strategy as outlined above.

### *Brucella* axenic growth

The concentration of *Brucella* cells (WT 2308 and Δ*bab2_0315*) was adjusted to 20% transmittance at OD_600_ and diluted into glucose-containing *Brucella* broth (Difco) at 1:20. MTT (3-(4.5-dimethylthiazol-2-yl)-2.5-diphenyltetrazolium bromide) reagent was added to the media to a final concentration of 0.5 mg mL^−1^, and cells were continuously incubated at 37°C in a Tecan plate reader, taking absorbance readings at 570 nm every 10 min for 72 h.

### *Brucella* axenic growth: compound cytotoxicity

NHI-2 (Sigma-Aldrich), 3-BPA, and 2-dG (Santa Cruz Biotechnology) were freshly prepared and serially diluted to test concentrations in a half-area 96-well plate in a total volume of 50 µL of PM9 medium (2 mM MgCl_2_• 6H_2_0, 1 mM CaCl_2_• 2H_2_O, 0.005% yeast extract, 0.005% Tween 40, 2.5 mM D-glucose, 5 mM sodium pyruvate, and 1X Dye Mix G [Biolog]). WT *Brucella abortus* strain 2308 was prepared from a plate culture in PM9 medium at 20% transmission at 600 nm, and 50 µL was added to each well for a total volume of 100 µL per well. Cells were incubated at 37°C, and absorbance at 630 nm was read every 2 h for 72 h using a Tecan microplate reader.

### XTT cell viability assay

The cytotoxicity of NHI-2, 3-BPA, and 2-dG to THP-1 cells was assessed using the XTT cell proliferation and viability assay kit (Roche), according to the supplier’s protocol. Briefly, cells were seeded at 50,000 cells/well. Various concentrations of compounds were added to each well, in triplicate, in the presence of 25 µg mL^−1^ of gentamicin. Cells were incubated for 72 h before measuring viability using the XTT assay. XTT reagents were prepared according to the manufacturer’s instructions. Following incubation, 50 µL of XTT labeling mixture was added per well, and cells were further incubated for 4 h at 37°C and 5% CO_2_. Absorbance measurements were collected at 590 nm and 750 nm (reference wavelength). Final data were expressed as absorbance readings at 590 nm (minus 750 nm reference absorbance) at the respective compound concentrations.

### Quantifying *Brucella* infection of THP-1 cells

THP-1 cells were seeded in a black-well, clear-bottom 96-well plate (Costar) at 50,000 cells/well, differentiated with 40 ng mL^−1^ PMA for 72 h. Two hours before infection, cells were treated with compounds, and infected with *B. abortus* at 100 MOI, as described above. Fresh compounds were added back into media containing 25 µg mL^−1^ gentamicin, and cells were further incubated for 72 h. After incubation, cells were washed three times with 1X PBS, and fixed for 10 min with 4% paraformaldehyde, followed by three washes with 1X PBS. Cell nuclei were labeled with 1 μg mL^−1^ Hoechst stain. Cells were imaged using a Leica-DMI6000B fluorescence microscope equipped with a 20X/0.4 NA objective, Hamamatsu Orca-R2 camera, and Leica Application Suite X. Forty images were captured in four replicate experiments: ten images per condition per replicate. Cell Profiler (v2.1.1) was used to measure the number of intracellular fluorescent Brucellae, and the number of THP-1 nuclei. The effect of the compounds on *B. abortus* intracellular growth was assessed by calculating the number of bacteria per cell, represented as a ratio of the total number of bacteria to the total number of nuclei (*Brucella* per THP-1 nucleus) per image, which was calculated as an average of 40 images ± standard error of the mean. Statistical significance was determined using one-way analysis of variance (ANOVA) followed by Dunnett’s test.

To enumerate CFU at 1, 24, and 48 hours post-infection of THP-1, *B. abortus* cells were resuspended in warmed RPMI and added at a multiplicity of infection (MOI) of 100 CFU per macrophage cell. To synchronize infections after the addition of *Brucella* cells, plates were centrifuged at 200 x *g* for 5 minutes. After a 30-minute incubation, the medium was removed and replaced with RPMI supplemented with 100 μg mL^−1^ gentamycin to kill extracellular bacteria. Afterwards, samples were incubated for 1 hour, after which monolayers were washed with PBS and THP-1 cells were lysed by addition of 0.01% Triton X-100. Serial dilutions were prepared and viable cells were determined by plating on TSA plates. For longer time points, the RPMI was removed, and replaced with RPMI supplemented with 50 μg mL^−1^ gentamycin. Additional samples were lysed as above and serial dilutions plated. All THP-1 assays were performed in triplicate utilizing independent *B. abortus* cultures.

## ACKNOWLEDGEMENTS

We thank the support staff of the Howard Taylor Ricketts Regional Biocontainment Laboratory. We also thank Barry Bochner, Aretha Fiebig, and Jennifer Mach for thoughtful feedback and critical evaluation of the manuscript. This study was funded in part by National Institutes of Health grants R01AI107159 and U19AI107792 to SC, and the Chicago Biomedical Consortium with support from the Searle Funds at the Chicago Community Trust.

## AUTHOR CONTRIBUTIONS

DC and SC conceived and designed the experiments, analyzed the data, and wrote the manuscript. DC and JW performed the experiments. All authors reviewed and approved the final manuscript.

